# The respiratory supercomplex from *C. glutamicum*

**DOI:** 10.1101/2021.06.16.448340

**Authors:** Agnes Moe, Terezia Kovalova, Sylwia Król, David J. Yanofsky, Michael Bott, Dan Sjöstrand, John L. Rubinstein, Martin Högbom, Peter Brzezinski

## Abstract

*Corynebacterium glutamicum* is a preferentially aerobic Gram-positive bacterium belonging to the Actinobacteria phylum, which also includes the pathogen *Mycobacterium tuberculosis*. In the respiratory chain of these bacteria, complexes III (CIII) and IV (CIV) form a CIII_2_CIV_2_ supercomplex that catalyzes oxidation of menaquinol and reduction of dioxygen to water. Electron transfer within the CIII_2_CIV_2_ supercomplex is linked to transmembrane proton translocation, which maintains an electrochemical proton gradient that drives ATP synthesis and transport processes. We isolated the *C. glutamicum* supercomplex and used cryo-EM to determine its structure at 2.9 Å resolution. The structure shows a central CIII_2_ dimer flanked by a CIV on each side. One menaquinone is bound in each of the Q_N_ and Q_P_ sites in each CIII, near the cytoplasmic and periplasmic sides, respectively. In addition, we identified a menaquinone positioned ~14 Å from heme *b*_L_ on the periplasmic side. A di-heme cyt. *cc* subunit provides an electronic connection between each CIII monomer and the adjacent CIV. In CIII_2_, the Rieske iron-sulfur (FeS) proteins are positioned with the iron near heme *b*_L_. Multiple subunits interact to form a convoluted sub-structure at the cytoplasmic side of the supercomplex, which defines a novel path that conducts protons into CIV.

## Introduction

In the final steps of energy conversion in aerobic organisms, electrons are transferred through the respiratory chain, which consists of membrane-bound proteins that transfer electrons from electron donors, such as NADH, to the final electron acceptor, O_2_. This electron current drives proton translocation from the negative (*n*) to the positive (*p*) side of the membrane, thereby maintaining a voltage difference and a proton concentration gradient that together generate a transmembrane proton motive force (PMF). The free energy stored in the PMF is used by the ATP synthase for production of ATP from ADP and phosphate, or to drive transmembrane transport (1).

NADH dehydrogenases and other enzymes transfer electrons to membrane-soluble quinone (Q), reducing it to quinol (QH_2_). In mitochondria and many bacteria, the reduced QH_2_ donates electrons to the cyt. *bc*_1_ complex, also known as complex III, which is found as an obligate dimer (CIII_2_). In each monomer of CIII_2_ the QH_2_ binds at the Q_P_ site, near the *p* side of the membrane. Energy conservation is realized through a bifurcated electron transfer from QH_2_, referred to as the Q-cycle (2) (**Figure 1**). The first electron from QH_2_ is transferred along the so-called C branch to a Rieske iron-sulfur protein, which harbors a redox-active 2Fe-2S (FeS) center. Oxidation of QH_2_ leads to the release of two protons to the *p* side of the membrane. The second electron is then transferred along the B branch, passing electrons sequentially to the low-potential heme *b*_L_ and the high-potential heme *b*_H_ before reducing Q bound in a second site, the Q_N_ site, located near the *n* side of the membrane. In canonical CIII_2_ the FeS center, which is bound in a mobile ectodomain, receives the electron from QH_2_ while in its B position in proximity to heme *b*_L_. Upon reduction of the FeS center and heme *b*_L_, the mobile FeS domain rotates by ~60° toward the *p* side to adopt its C position near cyt. *c*_1_. In the C position the electron from FeS is transferred first to heme *c*_1_ and then to a water-soluble cyt. *c* (3) (**Figure 1**). Cyt. *c* donates electrons to the last component of the respiratory chain, cytochrome *c* oxidase (also known as cyt. *aa*_3_ or complex IV, CIV). After exchange of Q for QH_2_ at the Q_P_ site of CIII, the sequence of events is repeated, resulting in formation of QH_2_ at the Q_N_ site and abstraction of two protons from the *n* side of the membrane, contributing further to the PMF (for review, see (4-9)).

**Figure 1.**
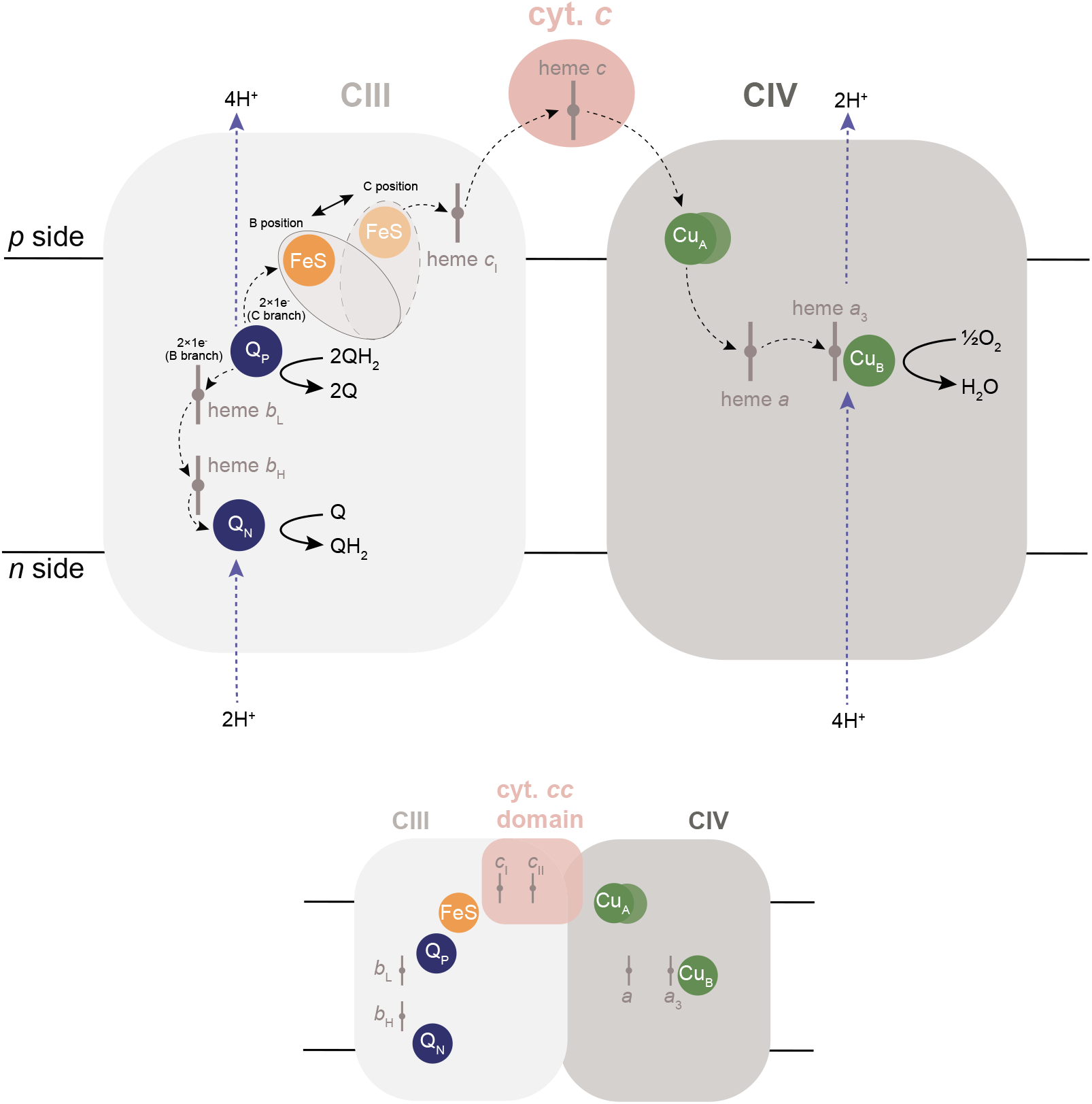
Reactions catalyzed by CIII and CIV. Upon binding of QH_2_ at the Q_P_ site in CIII one electron is transferred to FeS, along the C branch, and one along the B branch, via hemes *b*_L_ and *Q*_H_ to the Q at the Q_N_ site. The reaction sequence is repeated upon binding of a second QH_2_ at the Q_P_ site. In canonical CIII the mobile FeS ectodomain moves from the electron-receiving B position to the electron-donating C position to transfer an electron to cyt. *c*_1_, which reduced water-soluble cyt. *c*. Oxidation of QH_2_ at the Q_P_ site is associated with proton release to the *p* side of the membrane while reduction of Q at the Q_N_ site is associated with proton uptake from the *n* side. Reduced cyt. *c* transfers electrons to Cu_A_, heme *a*, and the heme *a*_3_/CuB catalytic site of CIV where O_2_ is reduced to H_2_O. Each electron transfer from cyt. *c* to the catalytic site is linked to pumping of one proton across the membrane. The lower part of the figure shows the *M. smegmatis* III_2_IV_2_ supercomplex where electrons from CIII to CIV are transferred via a di-heme cyt. *cc* domain.

The primary electron acceptor from cyt. *c* in CIV is a di-nuclear copper A site, Cu_A_, on the *p* side of the membrane. This copper center transfers electrons to heme *a* and then to the bi-nuclear catalytic site, composed of a heme *a*_3_ and copper Cu_B_. Upon electron transfer to the catalytic site, heme *a*_3_ binds an O_2_ molecule, which is reduced to H_2_O, in a process linked to proton uptake from the *n* side of the membrane (**Figure 1**). The free energy released upon oxidation of cyt. *c* and reduction of O_2_ is conserved by proton pumping from the *n* to the *p* side of the membrane (for review, see (10-12)).

Three major CIV families have been defined on the basis of amino-acid sequences as well as functionally important structural features such as proton pathways (13-15). Class A1 members are characterized by a XGHPEVY motif in subunit I and comprises the mitochondrial as well as a large number of bacterial CIVs, including the enzyme from *C. glutamicum*. In the XGHPEVY motif, H is a ligand of Cu_B_ (His265, *C. glutamicum* CIV numbering) while Y (Tyr269) is covalently linked to His265 (16-19). The A-type CIVs harbor two proton pathways, denoted D and K, used for proton uptake from the *n* side of the membrane (16-19). The K pathway transfers two protons to the catalytic site upon reduction of heme *a*_3_ and Cu_B_, while the D pathway is used for transfer of two protons to the catalytic site after binding of O_2_ to heme *a*_3_ and for all protons that are pumped across the membrane (20-24).

The respiratory chains of Gram-positive bacteria of the phylum Actinobacteria do not harbor genes for a water-soluble cyt. *c* (25) and the series of events that allows a Q-cycle is not as well understood. In these organisms CIII_2_ and CIV form an obligate CIII_2_CIV_2_ supercomplex in which electron transfer between CIII and CIV is mediated by a di-heme cyt. *cc* domain that replaces both cyt. *c*_1_ of the canonical CIII_2_ and the water-soluble cyt. *c*, as shown e.g. in *Mycobacterium smegmatis* (26-28) and *Corynebacterium glutamicum* (29-33) (**Figure 1**, lower part).

In order to gain insight into electron transfer and proton translocation in the CIII_2_CIV_2_ supercomplex we determined a high-resolution cryo-EM structure of the supercomplex from *C. glutamicum*. The structure shows density for a menaquinone (MQ) bound in each of the Q_N_ and Q_P_ sites of each CIII monomer. In addition, an MQ was found in a novel site on the membrane *p* side, ~14 Å from heme *b*_L_. As with the *M. smegmatis* supercomplex (26), an extended loop of the QcrB subunit covers the cytoplasmic opening of the D proton pathway of CIV, defining a novel proton-entry route via protonatable residues of QcrB. The FeS ectodomain in CIII_2_ was found to be locked in the B position, which suggests a Q-cycle mechanism that is gated only by local proton transfer rather than by FeS movement.

## Results and Discussion

### Isolation of the supercomplex

The 750 kDa *C. glutamicum* CIII_2_CIV_2_ supercomplex was purified using a Strep-tag on the CtaD subunit of CIV (29) (**supplementary Figure S1** and **Table S1**). Absorbance difference spectroscopy suggests a heme *a:b:c* ratio of approximately 1:1:1, consistent with the CIII_2_CIV_2_ composition of the supercomplex (29, 33). Mass spectrometry identified all of the subunits of CIII (QcrA, QcrB, QcrC) and CIV (CtaC, CtaD, CtaE), except for CtaF, which is a hydrophobic membrane protein that is likely difficult to detect by mass spectrometry, as suggested for the equivalent subunit in *M. smegmatis* (26). Three additional peptides associated with the CIII_2_CIV_2_ supercomplex were identified by mass spectrometry: P20 (later renamed to PRSAF1), P24, and P29 (later renamed to LpqE) (29). Peptide P24 was not identified in the structure. The MQH_2_:O_2_ oxidoreductase activity of the supercomplex was ~100 s^-1^, consistent with earlier measurements (29, 32).

### Overall structure of the supercomplex

To understand the mechanism by which the *C. glutamicum* III_2_IV_2_ supercomplex links electron transfer to proton translocation, we determined its structure by cryo-EM to a nominal resolution of 2.9 Å (**supplementary Figures S2-S3** and **Table S2**). Identifiers for proteins found in the *C. glutamicum* respiratory supercomplex are summarized in **Table S1**. The map shows that the core of the supercomplex is composed of a CIII_2_ dimer flanked by two distal CIV monomers (**Figure 2A**). This overall arrangement and its geometry is the same as the *M. smegmatis* supercomplex (26, 27).The core of CIV is composed of four subunits, CtaC-F (**Figure 2B**), while each protomer of CIII_2_ is composed of three subunits, QcrA-C (**Figure 2C**) (25, 33). An additional six subunits were identified in the *C. glutamicum* supercomplex structure (**Figure 2A**), two of which (LpqE and PRSAF1) were also found in the *M. smegmatis* supercomplex structure (27). We propose a new unifying nomenclature and refer to these additional subunits as AscX (Actinobacterial supercomplex, subunit X), except for LpqE (P29), which is an established name (where applicable, the previously-used names are given below in parentheses).

**Figure 2.**
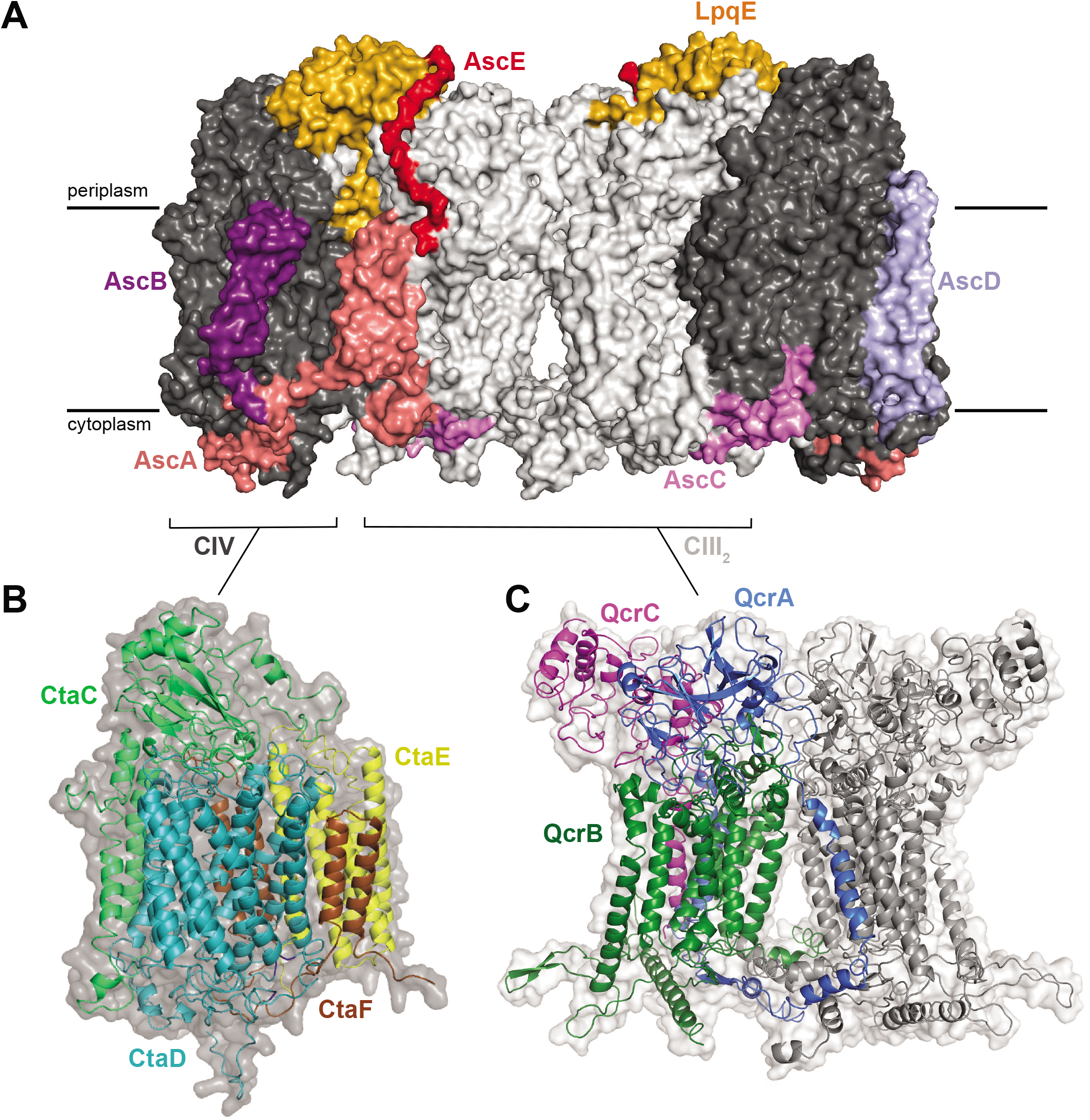
Structure of the supercomplex. (**A**) The entire III_2_IV_2_ supercomplex with core subunits in light (CIII_2_) or dark (CIV) grey, and accessory subunits in color as indicated. (**B**) The core of complex IV with CtaD, CtaC, CtaE and CtaF, shown in color and a surface representation of the entire CIV in dark grey. (**C**) The complex III2 dimer in which each monomer is composed of subunits QcrA-C. The right-hand side monomer is shown in grey while the subunits of the lefthand side monomer are colored. Note that TMH 1 of QcrA is integrated into the other half of the dimer. A surface representation of the entire CIII_2_ is shown in light grey.

The map allowed for construction of an atomic model for all subunits except for AscD and AscE (**supplementary Figure S4** and **Table S2**). On the periplasmic side of the supercomplex subunit LpqE is attached to the membrane via an N-terminal lipid anchor (**Figure 2A,** colored in gold). It also interacts with the cyt. *cc* domain of QcrC, and subunits QcrA, CtaC, and CtaD. As shown previously, LpqE did not co-purify with CIII und CIV alone, suggesting that the interaction with QcrC, which is absent in both single complexes, is necessary for the presence of LpqE. Consequently, LpqE may be involved in assembly of the supercomplex (29). Subunit AscA (colored in light salmon) is attached to both CIII_2_ and CIV. The N-terminal part of the protein starts with a short loop that continues to form two transmembrane *α*-helices attached to subunit QcrB of CIII_2_. A loop formed by 54 residues near the C terminus of AscA interacts with a loop of CtaD, and the two transmembrane *α*-helices. Like LpqE, AscA was not co-purified with the isolated CIII and CIV complexes, indicating that interaction with both QcrB and CtaD is required for co-purification.

A ~63-residue stretch of an unknown protein, denoted AscB (colored purple), was identified in the map. A tentative sequence was modeled based on the density and used to search the NCBI (ncbi.nlm.nih.gov) database for potential matches. This search identified protein GenBank: Cg0775, which provides a convincing representation of the complete density (**supplementary Figure S4B**). No protein from the mass spectrometric analysis provided a plausible match. AscB is folded into a transmembrane *α*-helical hairpin that is attached to CtaD. A short peripheral protein chain composed of 63 amino acid residues, named AscC (colored in violet) and also conserved in mycobacteria, was also modelled, as described above for AscB, based on GenBank: Cg0935 (**supplementary Figure S4B**). This protein is partially attached to QcrB on the cytoplasmic side of the supercomplex. The protein chain extends to contact CtaE and CtaF, as well as a QcrB loop that covers one of the proton pathways of CIV (see below). AscD (colored in blue) forms a transmembrane *α*-helical hairpin that is a part of CIV. It is located between the membrane-facing CtaF *α*-helical hairpin and CtaC *α*-helical hairpin. The sequence of this protein chain is unknown. Its N- and C-terminal loops are also part of a cytoplasmic side sub-structure, discussed in more detail below. AscE (colored in dark red) is attached to the FeS domain of QcrA and to LpqE. The resolution of this part of the structure is low and it was therefore modelled as polyalanine. The density also shows a similar lipid anchor to that of LpqE.

The structure of the *M. smegmatis* supercomplex showed a SodC-type Cu-containing superoxide dismutase (SOD) dimer, which is also composed of a lipobox motif attached to a lipoprotein segment (26, 27). The *C. glutamicum* strain used in the current study harbors only an Mn-containing SodA (34). AscE of the *C. glutamicum* supercomplex is located at the equivalent position of the *M. smegmatis* SodC N-terminal anchor (**Figure 2A** and **supplementary Figure S5**) and no SOD was found attached to the *C. glutamicum* supercomplex.

Density attributed to putative integral lipid molecules (35) was identified at 61 positions within the supercomplex. Unidentifiable lipids were modeled as hydrocarbon chains (**supplementary Figure S6**). Cardiolipin (CL), commonly found in membranes that are involved in maintaining an electrochemical proton gradient (36, 37), is identified at 14 positions. Four CL molecules are found at the interface of CIII and CIV in each half of the CIII_2_CIV_2_ supercomplex, consistent with a role in supporting supramolecular interactions in respiratory supercomplexes (38). A CL is also found at the monomer-monomer interface of the CIII_2_ dimer and one CL is bound to CtaD facing AscA and QcrB, further suggesting a role in higher-order assembly of complexes. In addition, another CL is found in a cavity defined by subunits CtaE and CtaF of CIV suggested to be used for O_2_ diffusion to the catalytic site (39, 40) (see below). All CLs are oriented with their negatively charged headgroups toward the *n* side of the membrane (see (36)).

### Overall structure of Complex III_2_

The FeS-containing ectodomain on the periplasmic side of QcrA is anchored by three transmembrane *α*-helices (TMH 1-3), one of which (TMH 1) is swapped between CIII monomers in the dimer and occupies the same position where the single transmembrane *α*-helix of the Rieske iron-sulfur protein is found in the canonical CIII (**Figure 2C**, colored in blue). The two additional transmembrane *α*-helices (TMH2 and 3) from QcrA are formed by an ~80 residue N-terminal extension not found in canonical CIII. TMH2 occupies the position where the transmembrane *α*-helix of subunit cyt. *c*_1_ is found canonical CIII, which in the *C. glutamicum* structure is shifted towards the middle of the supercomplex (see also (26)). The FeS ectodomain of the *C. glutamicum* CIII is fixed in the B position by the LpqE subunit on the periplasmic side of the protein, which was also noted in one structure of the *M. smegmatis* supercomplex (27). This tight interface between the QcrA ectodomain and cyt. *cc* would preclude movement of the ectodomain.

The QcrC subunit is composed of a di-heme cyt. *cc* (cyt. *c*_I_ and *c*_II_, **Figure 1**, lower part) head domain and a transmembrane *α*-helix, which is displaced compared to that of cyt. *c*_1_ subunit in the canonical CIII (**Figure 2C**, colored in magenta). The C-terminal sequence of QcrC forms a single transmembrane *α*-helix that contacts QcrB. Cyt. *c*_I_ of the cyt. *cc* domain interacts with the QcrA ectodomain on the opposite side from the FeS center, while cyt. *c*_II_ is bound near the electron-accepting Cu_A_ site of CIV. This arrangement of cyt. *c*_I_, cyt. *c*_II_, and Cu_A_ provides an electronic connection between CIII and each CIV of the supercomplex.

Subunit QcrB (**Figure 2C**, colored in green) consists of 8 transmembrane *α*-helices and harbors hemes *b*_L_ and *b*_H_, which occupy the same positions as in the canonical (41) and *M. smegmatis* (26, 27) CIII_2_. In addition, the Q_P_ and Q_N_ quinone-binding sites are defined in part by residues of QcrB. The C terminus of QcrB is extended by 137 residues, not present in the canonical CIII, on the cytoplasmic side of the supercomplex (30). About 20 of these residues form a loop that contacts the CIV subunit CtaD (26).

### Quinone binding in complex III

The Q_P_ site is typically empty in X-ray crystal structures of canonical CIII_2_ from a wide range of organisms, and was identified from the positions of the inhibitors stigmatellin and myxothiazol in inhibitor-bound structures of the complex (6). A recent cryo-EM study revealed ubiquinone (UQ) bound at the Q_P_ site of the CI-CIII_2_ mammalian supercomplex, but only in one monomer of CIII_2_ (42). The map of CIII_2_CIV_2_ supercomplex reveals density for MQ adjacent to the FeS cluster in each CIII monomer, thereby defining the Q_P_ site in *C. glutamicum* (**Figure 3A,B** and **supplementary Figure S7A**). This site overlaps with the UQ site identified in mammalian CIII (42) (**supplementary Figure S7B**). We designate the MQ molecule in this position as MQ1a (**Figure 3A-C**). In structures of the *M. smegmatis* supercomplex density was seen at a distal position near the entrance to the Q_P_ cavity (26, 27), which we designated as MQ1b (**supplementary Figure S7C**).

**Figure 3.**
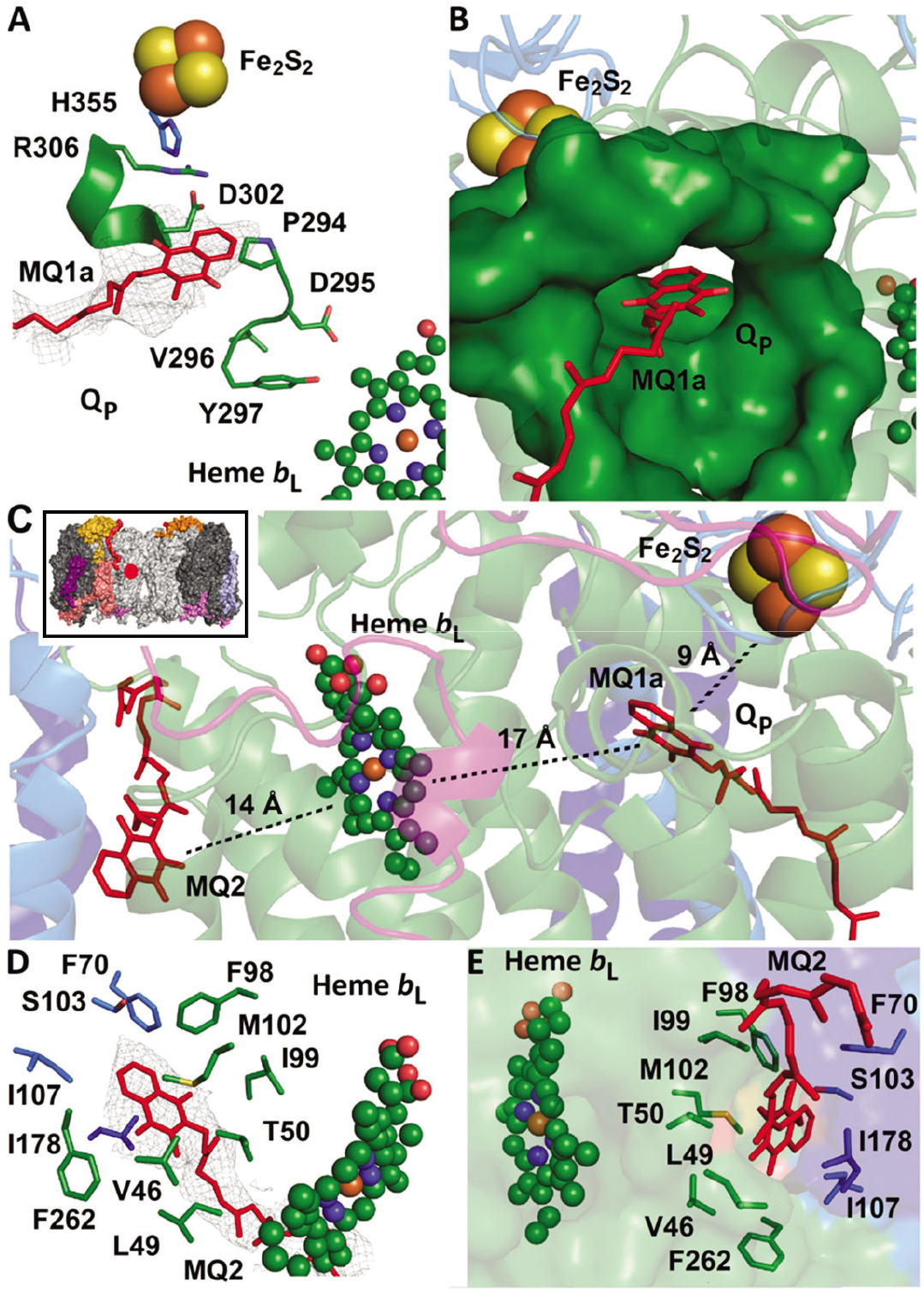
Q-binding sites on the periplasmic side. (**A**) Density modeled as an MQ in the Q_P_ site, MQ1a. Protonatable residues discussed in the text and the PDVY motif in the QcrB subunit are shown. Residue H355 is found in subunit QcrA. (**B**) The Q_P_ site with bound MQ1a. (**C**) Position of MQ1a in relation to the FeS cluster and heme *b*_L_, as well as a second MQ in site MQ2. In the inset the approximate position of this area in one CIII monomer of the supercomplex is marked with a red circle. (**D,E**) Amino-acid residues that define the MQ2 binding site. The MQ2 density is shown in panel **D**.

The Q_P_ cavity in the *C. glutamicum* CIII is larger than in the *M. smegmatis* enzyme. The density corresponding to MQ observed in the Q_P_ site of the *C. glutamicum* CIII is diffuse, spanning across a position equivalent to MQ1b (**supplementary Figure S7A**). This diffuse density could correspond to a single MQ bound only at MQ1a, or to averaging of two CIII populations in which one MQ bound in either MQ1a or MQ1b. In either case, observation of MQ at positions MQ1a or MQ1b in *C. glutamicum* and *M. smegmatis*, respectively, suggests that there are two possible binding modes in or just outside of the Q_P_ site, respectively. Two QH_2_ binding positions were suggested in a proposed mechanism for canonical CIII_2_ in which QH_2_ initially binds in a “stand-by” site and is then re-located into an oxidation site that is formed transiently after docking of the FeS domain in the B position of (7). Furthermore, early EPR data indicated two Q-binding positions in the Q_P_ site of *R. capsulatus* cyt. *bc*_1_ (43). Two possible MQ binding positions inside and just outside of the Q_P_ cavity is also reminiscent of the two Q-binding positions near the Q_B_ site in a crystal structure of a dark-adapted photosynthetic reaction center from *Rhodobacter sphaeroides* (44).

Structural studies of the *M. smegmatis* CIII_2_CIV_2_ supercomplex (26) identified density corresponding to MQ in an unexpected location, near the Tyr of the PDFY motif in CIII, which is equivalent to the canonical PEWY Q-binding motif (45). This MQ is at the vertex of a triangle formed with heme *b*_L_ (~20 Å) and the FeS center (~20 Å) (**supplementary Figure S8A**). In the *C. glutamicum* structure the corresponding position in the enzyme cannot accommodate MQ because it is occupied by Trp265 (**supplementary Figure S8B**). The positions of Trp265 in *C. glutamicum* and the equivalent Trp276 in *M. smegmatis* differ, presumably because the former accommodates a smaller Val instead of Phe in its Q-binding motif (PDVY in *C. glutamicum*) (**supplementary Figure S8BC**). In *C. glutamicum* we observed MQ, which we designate MQ2 at a different location at the *p* side of the membrane, 14 Å from heme *b*_L_ and 30-35 Å from the Q_P_ site (**Figure 3C-E** and **supplementary Figure S8A**). The equivalent location in the *M. smegmatis* structures harbors a lipid tail density (26, 27). Identification of a second Q-binding site on the *p* side of CIII in both *C. glutamicum* and *M. smegmatis* suggests a functional role, which is discussed below.

MQ is also found in the Q_N_ site (**supplementary Figure S9** colored in pink), with the head group at the same position seen previously in the canonical (41) and *M. smegmatis* (26, 27) CIII.

### The Q-cycle of complex III

The bifurcated electron transfer from QH_2_ at the Q_P_ site is fundamental for the Q-cycle mechanism that conserves energy in CIII (**Figure 1**) (46). As outlined in the Introduction section, in this process the first electron from QH_2_ is transferred to FeS while the second is transferred to heme *b*_L_:

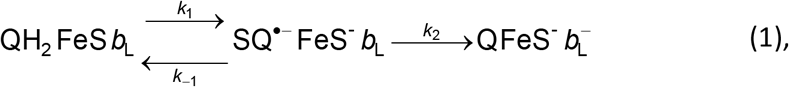

where 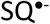 is a putative semiquinone. In canonical CIII, rotation of the FeS domain from the electron-receiving B position to the electron-donating C position near cyt. *c*_1_ is thought to be part of the mechanism that allows bifurcated electron transfer (6, 7, 9). However, this movement of the FeS domain is not required for electron bifurcation (47).

Oxidation of QH_2_ at the Q_P_ site has been studied primarily with the canonical CIII (7, 48, 49). Upon QH_2_ binding, the FeS domain moves to the B position and QH_2_ forms a hydrogen bond with the FeS ligand His181 (*S. cerevisiae* numbering), which also receives a proton from QH_2_ upon electron transfer to FeS (50-54). The second electron is transferred to heme *b*_L_, along with proton transfer, presumably to Glu272, which is part of the Q-binding PEWY motif in *S. cerevisiae* (7, 49, 51). The protonated Glu272 is suggested to rotate toward the heme *b*_L_ propionate, which transiently binds the proton (49, 51) before it is released to the *p* side aqueous phase. After transfer of the second electron to heme *b*_L_ the FeS domain moves to the C position, which allows electron transfer to cyt. *c*_1_ and release of the proton from His181 to the aqueous phase on the *p* side.

Because the *C. glutamicum* FeS domain is locked in the B position, a mechanism other than movement of the FeS domain must exist to explain the proton-coupled electron-transfer reactions linked to MQH_2_ oxidation. Similar to the canonical CIII, it is feasible that in the *C. glutamicum* enzyme an electron and a proton are transferred from MQH_2_ in the Q_P_ site to FeS and its His355 ligand (**Figure 3A**), respectively (**Figure 4**, e^-^_1_ and H^+^_1_, top left). The equivalent of *S. cerevisiae* Glu272 in *C. glutamicum* is Asp295 (**Figure 3A**) of the PDVY motif, which is located near the heme *b*_L_ propionates, but the shorter Asp side chain is too short to reach MQH_2_ in the Q_P_ site to accept a proton. Instead, Asp302 (**Figure 3A**), located ~5 Å from MQH_2_ in the Q_P_ site could receive a proton upon transfer of the second electron from 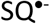 to heme *b*_L_ (**Figure 4**, e^-^_2_ and H^+^_2_, top left). In many Actinobacteria a Glu residue is found at this position, which could also serve as a proton acceptor. Arg306 is ~3 Å from Asp302 (**Figure 3A**), which points to a possible route for proton release to the *p* side aqueous solution (**Figure 4**, top right). Interestingly, Asp302 is ~3 Å from His355 suggesting that this residue is also on a proton-transfer pathway from His355. This architecture offers a plausible mechanism for Q-cycle electron branching in *C. glutamicum* that is guided by local proton-transfer reactions in the protein matrix. We suggest that after the initial oxidation of MQH_2_ (**Figure 4**, top left), the electron is stabilized by the His355 proton. Thus, electron transfer from FeS^-^ to heme *c*_I_ is not possible because His355 cannot be deprotonated until H^+^_2_ is released from Asp302. If electron transfer from heme *b*_L_ to heme *b*_H_ occurs as fast as proton transfer from Asp302 to the *p* side (**Figure 4**, top right), charge separation along the B branch is accomplished while FeS remains in the reduced state, FeS^-^. The FeS ligand His355 can become deprotonated only after Asp302 loses its proton, which allows electron transfer from FeS^-^ to heme *c*_I_, along the C branch (**Figure 4**, bottom right). In the final step of the reaction the second proton from D302 is released via R306 (**Figure 4**, bottom left). This model is supported by the observation that the FeS^-^ → heme *c*_I_ electron transfer is the slowest of the measured electron-transfer reactions in the *C. glutamicum* supercomplex (32).

**Figure 4.**
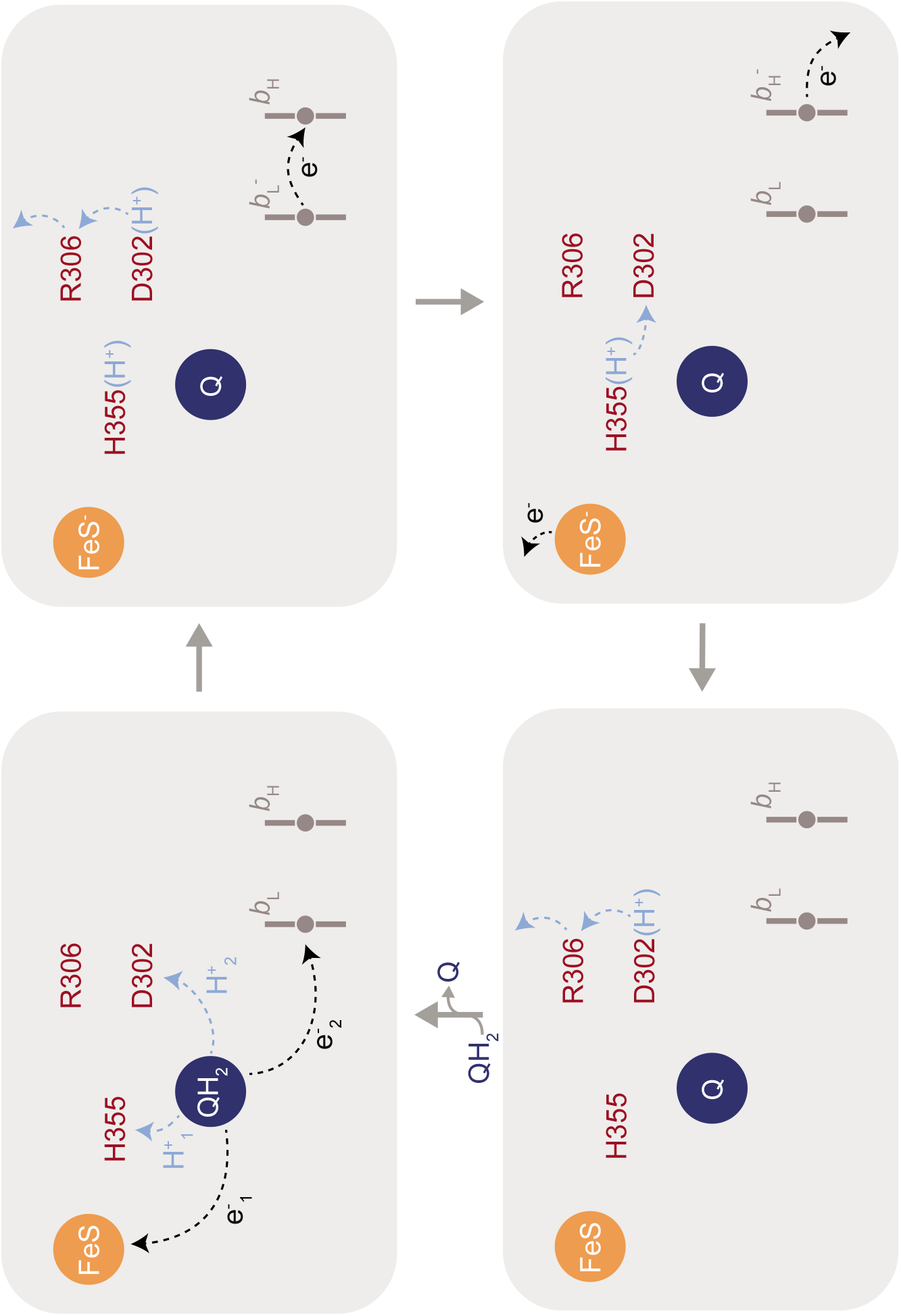
The Q-cycle electron bifurcation in *C. glutamicum* CIII. The first electron and proton from MQH_2_ are transferred to FeS and His355, respectively. The second electron and proton are transferred to heme *b*_L_ and Asp302, respectively. After this charge separation electron transfer from FeS^-^ (to heme *c*, not shwon) is not possible because His355 cannot be deprotonated until the proton is released from Asp302. This sequence of events ensures that the second electron is transferred along the B branch before the first electron is transferred along the C branch.

A second MQ is found at the *p* side of the supercomplex, 14 Å from heme *b*_L_ (MQ2 in **Figure 3C-E**). The position of MQ2 is different from that observed in the *M. smegmatis* enzyme (26, 27) (**supplementary Figure S8**). The role of this MQ is unknown, but we speculate that electron transfer via the MQ2 site could provide an alternative electron path that bypasses heme *b*_H_, thereby decoupling electron transfer through the CIII portion of the supercomplex from generation of a PMF and preventing energy conservation. This pathway would maintain an electron flux through the respiratory chain, for example at low O_2_ concentrations (1).

### Complex IV

The core subunits of *C. glutamicum* CIV, CtaC and CtaD, are homologous to conserved subunits (SU) II and I, respectively, of canonical CIV. These subunits harbor all redox-active metal cofactors of CIV. Subunit CtaC is composed of two transmembrane *α*-helices and a head domain, which binds the primary electron acceptor, Cu_A_. Subunit CtaD is composed of 12 transmembrane *α*-helices, which bind heme *a* and form the catalytic site that includes heme *a3* and Cu_B_. The relative positions of the redox-active cofactors within CtaC and CtaD are the same in *C. glutamicum* as in CIV from other organisms (**Figure 5A**). In canonical CIV the seven transmembrane *α*-helices of SU III form a V-shaped O_2_ channel (39, 40) that harbors three tightly-bound lipid molecules (55). As with the *M. smegmatis* supercomplex (26, 27), SU III from canonical CIV is replaced by two proteins, CtaE and CtaF, in the *C. glutamicum* enzyme (**Figure 2B**, brown and yellow) harboring a single CL molecule (**supplementary Figure S6**). The division of SU III into two parts resembles the supercomplex structure of alternative complex III (ACIII) and CIV in *Flavobacterium johnsoniae* where the equivalent of SU III has lost the first two transmembrane *α*-helices (equivalent of CtaF) (56).

**Figure 5.**
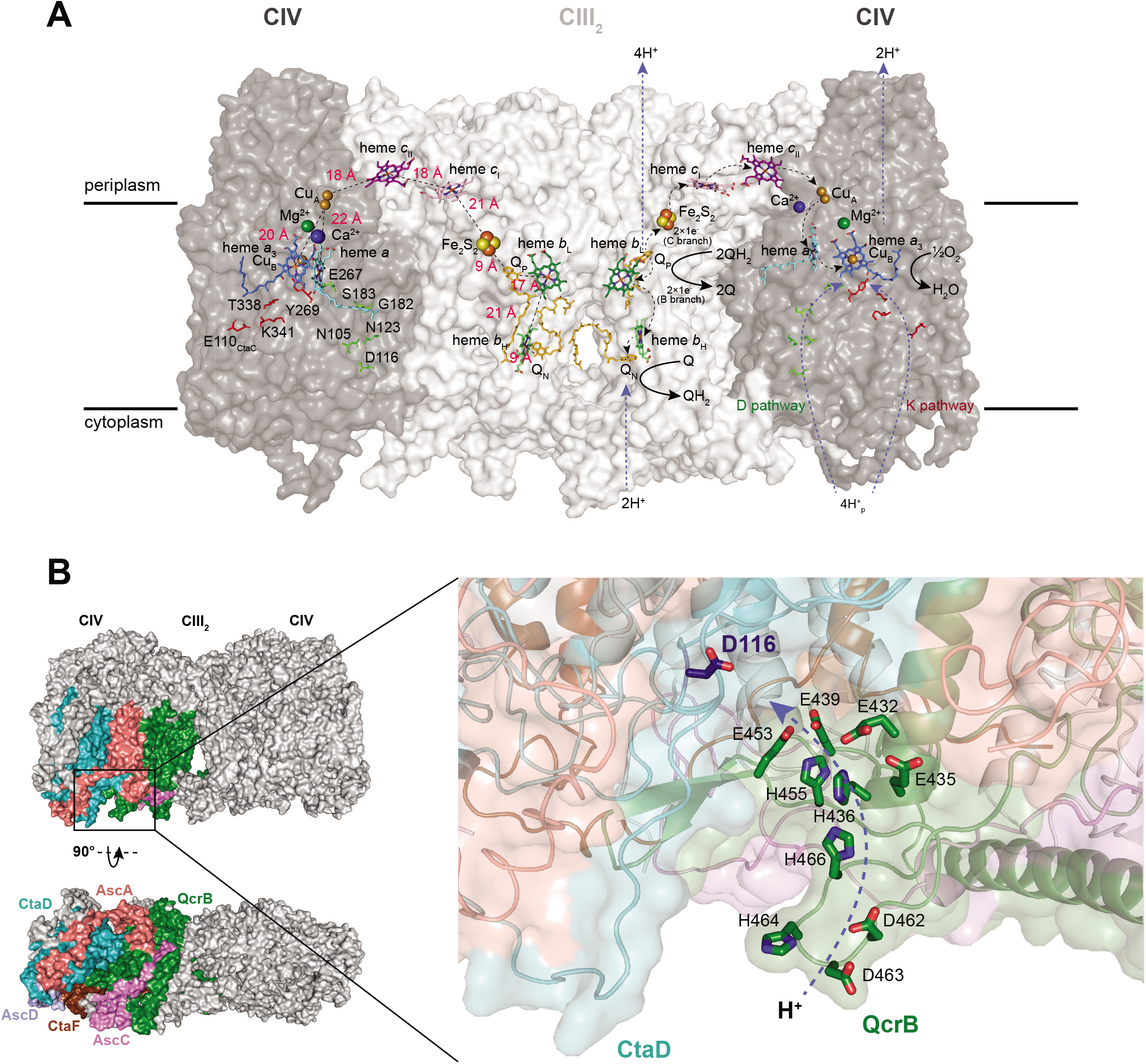
The *C. glutamicum* supercomplex. (**A**) The right- and left-hand side halves of the supercomplex show the reactions catalyzed by CIII and CIV, and distances between the cofactors, respectively. Residues of the D and K pathways in CIV are labelled in the left-hand side. (**B**) The entry point of the D pathway in complex IV. Subunits that contribute residues that define the entry point of the D pathway of CIV are shown in color. A complex III QcrB loop (green) forms a “lid” over the D-pathway entry point cavity. This “lid” harbors a number of protonatable residues that define a proton pathway leading from the cytoplasmic side to the D pathway entry point defined by Asp116.

Previous structures of mammalian, *S. cerevisiae*, and bacterial CIV revealed a Mg^2+^ ion 12 Å from Cu_B_ and 13 Å from the heme *a*_3_ iron (18, 57-59). In addition, a Na^+^ or Ca^2+^ ion was found to be bound in mitochondrial and bacterial CIV, respectively, at a specific site on the *p* side of CIV (18, 58, 59). In the present structure we found densities at both positions, which we modelled as Mg^2+^ and Ca^2+^, respectively (**Figure 5A**, green and blue spheres, respectively).

The supercomplex structure reveals both the K and D proton pathways in CIV. The K proton pathway, used for proton transfer to the catalytic site upon reduction of heme *a*_3_ and Cu_B_, starts near the *n*-side surface at Glu110 (CtaC), and is lined by a number of CtaD residues, including the conserved central Lys341 as well as Tyr269 at the catalytic site (21, 22, 60, 61). The D proton pathway is defined by the highly-conserved Asp116 in the inner part of a cavity in CtaD and a number of polar residues that span the distance to the highly-conserved Glu267 of the XGHPEVY motif (**Figure 5**). X-ray crystal structures of CIV from other organisms also revealed about ten water molecules that span the distance between Asp116 and Glu267 (16-19), but these water molecules could not be resolved in the current cryo-EM map. The D pathway is used for proton transfer to the catalytic site as well as for proton pumping to the *p* side of the membrane after binding of O_2_ to heme *a*_3_ (20, 22, 60, 61).

### Cytoplasmic side sub-structure

At the cytoplasmic side of the *C. glutamicum* supercomplex multiple subunits interact to create an intricate sub-structure, which includes secondary structure elements from CtaD, the C terminus of AscA, a CtaF loop, AscC, and the N and C termini of AscD (**Figure 5B**). The equivalent sub-structure of the *M. smegmatis* supercomplex is composed of fewer components, primarily subunits MSMEG_4692 (CtaI) and MSMEG_4693 (CtaJ) (26, 27) that are not present in the *C. glutamicum* supercomplex. Interestingly, as seen in the *M. smegmatis* supercomplex, the 20-residue QcrB loop (see above) covers the proton-uptake cavity around Asp116 (**Figure 5B**), which in the canonical CIV is exposed to the *n*-side aqueous solution. Conservation of this feature in both the *M. smegmatis* and *C. glutamicum* supercomplexes suggests a functional role. Mutation of the equivalent of Asp116 or residues in its vicinity in other CIVs result in drastic changes in the proton-uptake rate or uncoupling of proton pumping from O_2_ reduction (62-64). Furthermore, mutation of a Glu residue in a C-terminal flexible loop in *R. sphaeroides* CIV, 10 Å “below” the equivalent of Asp116, near the QcrB extension loop in the *C. glutamicum* structure, resulted in a decrease in the proton-pumping stoichiometry by a factor of two (65). Collectively, these data show that the area around the D pathway opening is critical for determining proton-uptake kinetics, presumably by tuning the electrostatic potential thereby providing a proton-collecting antenna (66). In addition, in *C. glutamicum* a chain of Asp, Glu, and His residues at the C terminus of QcrB provide an alternative proton path from the *n* side surface to Asp116 (**Figure 5B**).

We speculate that the structural modification of the cytoplasmic side sub-structure in *C. glutamicum* is result of differences between the cytoplasmic composition of Gram-negative alphaproteobacteria and mitochondria compared to Gram-positive Actinobacteria, as evident from a higher turgor pressure for the latter (67-69). As outlined above, the proton-collecting antenna around Asp116 determines the proton-uptake kinetics by the D pathway in CIV and any changes to the ionic composition of the cytoplasm are expected to modify this kinetics (66). We propose that the structural modifications around the D pathway opening in *C. glutamicum* optimize the proton-collecting function for the cellular environment of actinobacteria.

### Electron transfer from QH_2_ to O_2_

Complexes III and IV are electronically connected by the cyt. *cc* domain of the QcrC subunit of CIII. In the *C. glutamicum* structure the distance between FeS and cyt. *c*_I_ is 21 Å (**Figure 5A**). Assuming a Δ*G*^0^ ≅ +60 meV (33) and a reorganization energy of 0.7 eV yields a time constant for electron transfer from FeS to cyt. *c*_I_ of ~10 ms (70), which is consistent with the measured value of ~6.5 ms for oxidation of the *C. glutamicum* CIII (32). The Fe-Fe distances between hemes *c*_I_ and *c*_II_, and between heme *c*_II_ and Cu_A_ are both ~18 Å (**Figure 1**) yielding calculated time constants of ~1 ms for each electron transfer. Kinetic data showed biphasic oxidation of cyt. *cc* suggesting a time constant for electron transfer to Cu_A_ in the range 100 μs - 2 ms (32), which is consistent with the estimated value.

### Summary

This study reveals the structure of the respiratory CIII_2_CIV_2_ supercomplex from *C. glutamicum*. The structure shows MQ bound inside the Q_P_ site cavity and offers insights into how actinobacteria enable energy-conserving Q-cycle electron bifurcation without the mobile FeS domain found in canonical CIII. The structure shows a novel D proton pathway at the opening of CIV where residues from the neighboring QcrB subunit provide a proton-entry route from the *n* side. These findings illustrate the wide variety of structures that allow realization of respiratory pathways in aerobic organisms with particular insight into respiration in actinobacteria such as *M*. *tuberculosis* where respiration is a validated drug target.

## Materials and methods

### Growth of bacteria

*Corynebacterium glutamicum*, strain ΔC-D_st_ (*13032ΔctaD* with pJC1-*ctaD*_St_), described before (29), was grown on BHI-Agar plates (33 g/l brain heart infusion broth, 15 g/l agar, 20 g/l D-(+)-glucose, 25 mg/l kanamycin). Single colonies were picked, inoculated into 10 ml BHI culture medium and grown over night using a shaker at 300 rpm, 30°C. The pre-culture was diluted into 500 ml CGXII medium (71) in a 2 l flask and shaken at 160 rpm, 30°C until the OD_600_ reached 12. The cells were again diluted into 2 l CGXII medium in a 5 l baffled flask and shaken at 130 rpm, 30°C. The cells were harvested at OD_600_ 17, 10 000 x *g* for 30 min, JLA 8.1000 rotor (Beckman).

### Membrane preparation

Cells were homogenized in 4 ml cell lysis buffer (100 mM Tris-HCl at pH 7.5, 5 mM MgSO_4_) per gram of cells in the presence of a few crystals of a protease inhibitor phenylmethanesulfonyl fluoride (Sigma) and DNaseI (Roche). Cells were broken with a cell disrupter with 4 cycles at 35 kPsi (Constant Systems) and cellular debris was removed by centrifugation at 90 000 x *g* for 20 min at 4°C (45 Ti rotor, Beckman). Membranes were collected by ultracentrifugation at 220 000 x *g* for 90 min at 4°C (45Ti rotor, Beckman).

### Isolation of supercomplexes

Membranes were solubilized in 100 mM Tris-HCl pH 7.5, 100 mM NaCl, 2 mM MgSO_4_, 50 mg/l avidin (to prevent unspecific binding to the column, see below), 1% (w/v) DDM to a protein concentration of 5 mg/ml and incubated for 45 min, at 4°C under gentle stirring. Insolubilized material was removed by ultracentrifugation at 39 000 x *g*, 20 min, 4°C (SW41 rotor, Beckman). The supernatant was concentrated with a 100-kDa molecular weight cutoff concentrator (Merck Millipore) until the volume was reduced to 5 ml. The concentrated supernatant was then diluted in solubilization buffer without detergent to yield a final DDM concentration of 0.1 % (w/v) and concentrated again to reach a volume of <10 ml. The concentrated supernatant was applied to a gravity flow Strep-Tactin Superflow column (4 ml bed volume, Iba Lifescience). The column was washed 3 times with 0.5 column volumes of washing buffer (100 mM Tris-HCl pH 7.5, 100 mM NaCl, 2 mM MgSO_4_, 0.05% (w/v) DDM). The protein was then eluted with 3 column volumes of elution buffer (100 mM Tris-HCl pH 7.5, 100 mM NaCl, 2 mM MgSO_4_, 0.05 % (w/v) DDM, 2.5 mM D-desthiobiotin). The eluted supercomplex solution was concentrated as described above and further purified by size exclusion chromatography on a Superose 6 Increase 10/300 GL column (GE Healthcare), preequilibrated with buffer (100 mM Tris-HCl pH 7.5, 100 mM NaCl, 2 mM MgSO_4_, 0.05 % DDM) using an Äkta Pure M25 chromatography system (GE Healthcare) operated at 4°C with UV detection at 280 nm and 415 nm. Collected fractions containing supercomplex were concentrated for further analysis.

### Spectral analysis

The purified supercomplex was analyzed by UV–visible absorption spectroscopy (Cary 100 Spectrophotometer, Agilent Technologies). Difference spectra of the dithionite-reduced and oxidized states of the supercomplex were recorded. The reduced - oxidized difference absorption coefficients used to estimate the cofactor stoichiometry were: ≡_605–630_ = 24 mM^-1^cm^-1^ (cyt. *aa*_3_), ε_562–577_ = 22 mM^-1^ cm^-1^ (cyt. *b*), and ε_552-540_ = 19 mM^-1^ cm^-1^ (cyt. *c*) (28, 29).

### Preparation of menaquinol

2,3-dimethyl-[1,4]naphthoquinone (1.8 mg) was dissolved in 0.5 ml ethanol to yield a 20 mM solution. Several crystals of sodium borohydride (NaBH4) were added to reduce the quinone to quinol. The solution was kept on ice until transparent and HCl was added until formation of small bubbles in the solution ended. The sample was centrifuged for 10 min at 10 000 x *g* and the supernatant containing reduced quinol was aliquoted, flash frozen, and stored in −80°C until use.

### Activity assays

The O_2_ reduction rate was measured at 25°C using a Clark-type oxygen electrode in a buffer containing 100 mM Tris-HCl pH 7.5, 100 mM NaCl, 2 mM MgSO_4_, 0.05 % DDM. The reaction was initiated by addition of 5 μl of 20 mM 2,3-dimethyl-[1,4]naphthoquinol (Rare Chemicals GmbH) solution into a 1 ml chamber containing the supercomplex solution (40 nM). The activity was obtained from the initial slope of the graph. The background O_2_-reduction rate was measured and subtracted from the O_2_-reduction rate obtained in the presence of the supercomplex.

### Gel electrophoresis

Blue Native (BN) PAGE was performed according to the manufacturer’s instruction with pre-cast gel, NativePAGE™ 4-16 % Bis-Tris (Thermo Fisher Scientific). The gel was run at 4°C for 60 min at 150 V, then the cathode buffer was exchanged to anode buffer and run for an additional 40 min at 250 V. The gel was then stained with Coomassie Brilliant Blue. The band corresponding to supercomplex from BN PAGE was subjected to mass spectrometry, which indicated the presence of both complexes III2 and IV, as well as two additional subunits, LpqE and AscA (P20, PRSAF1). AscB and AscC were not identified in the mass spectrometric analysis.

SDS PAGE was performed according to manufacturer’s instruction with pre-cast gel, NuPAGE™ 4-12% Bis-Tris (Thermo Fisher Scientific) in MES running buffer. Samples were heated at 65°C for 30 min and run at 4°C for 45 min at 200 V. The gel was then stained with Coomassie Brilliant Blue.

### Grid preparation and cryo-electron microscopy

Purified supercomplexes (3 μl) at a concentration of 10 mg ml^-1^ was applied to holey carbon film coated copper EM grids (C flat 2/2 3C T50) that had been glow-discharged in air for 120 s at 20 mA (PELCO easiGlow). Grids were blotted for 3 s at 4°C and 100 % humidity before being plunge-frozen in liquid ethane with a Vitrobot Mark IV (Thermo Fisher Scientific). Cryo-EM images were collected at 300 kV with a Titan Krios electron microscope (Thermo Fisher Scientific) equipped with a Gatan K2-summit direct electron detector and a Bio-quantum energy filter (Gatan). Data were collected with a nominal magnification of 130 000 x, corresponding to a calibrated pixel size of 1.06 Å. Automated data collection was done with the EPU software package (Thermo Fisher Scientific). A dataset of 2768 movies was collected, each consisting of 40 exposure fractions. The camera exposure rate and the total exposure of the specimen were 7.7 e^-^/pixel/s and 55 e^-^/Å^2^, respectively (**Table S2**).

### Image analysis

All image analysis was performed with cryoSPARC v2 unless otherwise stated (72). Movies were aligned with *MotionCor2* (73) and contrast transfer function (CTF) parameters were estimated in patches. Templates for particle selection were generated by 2D classification of manually selected particle images. A total of ~560 000 particle images were selected, images were corrected for local motion (74) and extracted in 310 × 310 pixel boxes. The dataset was first cleaned with 2D classification and then with three rounds of *ab initio* 3D classification and heterogeneous refinement reducing the size to ~65 000 particle images. Local and global CTF refinement followed by homogeneous refinement without the application of symmetry resulted in a map at 2.9 Å resolution.

Figures were prepared using the software PyMOL (Molecular Graphics System, Version 2.0 Schrödinger, LLC., (75)) as well as UCSF Chimera, developed by the Resource for Biocomputing, Visualization, and Informatics at the University of California, San Francisco, with support from NIH P41-GM103311.

### Model building and refinement

An initial model of all the subunits of the *C. glutamicum* supercomplex was built manually into the C1 symmetry density map with 2.9 Å resolution using Coot (76). Subunits AscB, AscC and AscD were initially build as a poly-alanine chains. Sequences of AscB and AscC were then built into the density map where high enough resolution allowed identification of amino-acid side chains, based on known sequences of the *C. glutamicum* genome using BLAST (77). No sequence match was found for AscD; it remained modelled as a poly-alanine chain. The model was refined using combination of phenix_real_space_refine (78) and manual adjustments in Coot.

## Data deposition

Data deposition: all electron cryomicroscopy maps described in this article have been deposited in the Electron Microscopy Data Bank (EMDB) (accession nos. EMD-XXXX to EMD-XXXX).

## Acknowledgements

We thank Mikael Oliveberg for valuable discussions. This work was supported by the Knut and Wallenberg Foundation (MH, PB), the Swedish Research Council (MH, PB), and Canadian Institutes of Health Research grant PJT162186 (JLR). JLR was supported by the Canada Research Chairs program. Cryo-EM data was collected at the Swedish National Cryo-EM Facility funded by the Knut and Alice Wallenberg, Family Erling Persson and Kempe Foundations, SciLifeLab, Stockholm University and Umeå University. Mass spectrometry was done at the Mass Spectrometry-based Proteomics Facility at Uppsala University.

## Notes

### Competing Interest Statement

The authors have declared no competing interest.

