## Supplementary material for "The respiratory supercomplex from *C. glutamicum*"

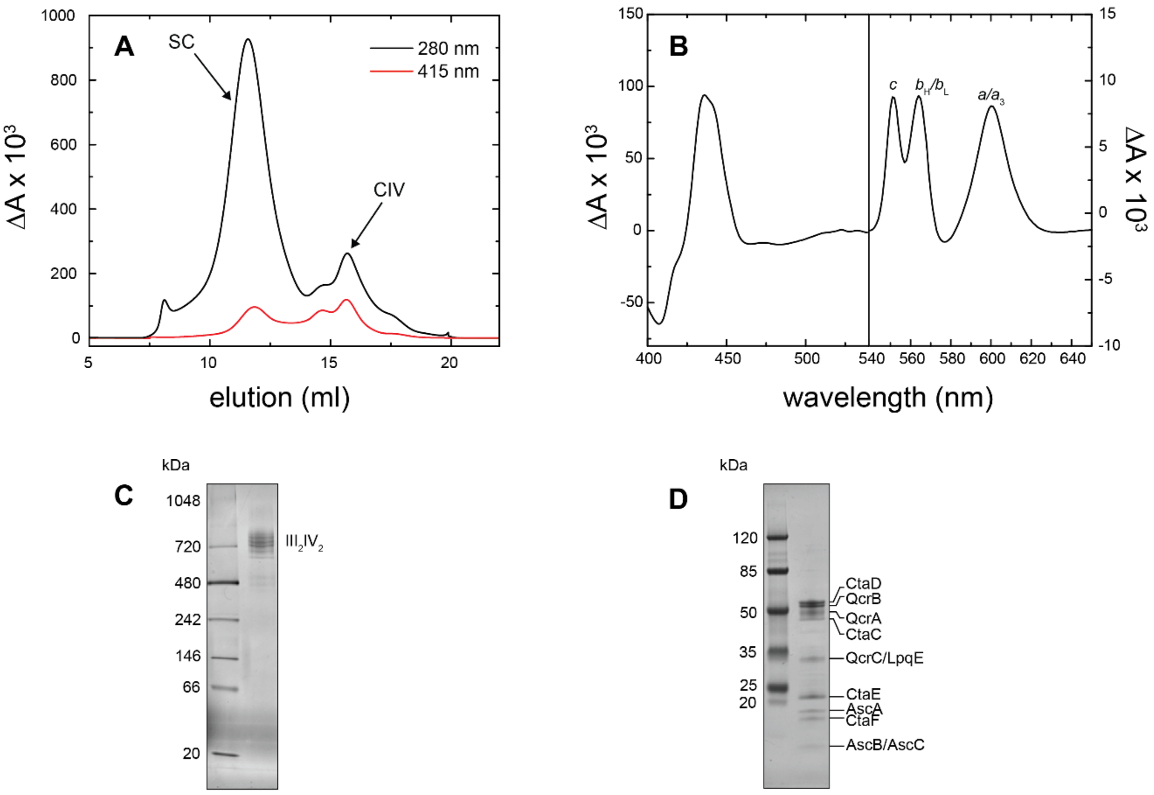

**Figure S1**. **Purification of the *C. glutamicum* $III_2IV_2$ supercomplex.** (**A**) Elution profile from size-exclusion chromatography with a Superose 6 Increase 10/300 GL column in 100 mM Tris-HCl pH 7.5, 100 mM NaCl, 2 mM $MgSO_4$, 0.05% DDM. The elution process was followed at 280 nm (total protein) and 415 nm (hemes). Dithionite reduced minus oxidized difference spectra of the fractions showed the presence of supercomplexes (SC), eluted in the first peak, and free CIV in the second peak. (**B**) Dithionite reduced-minus-oxidized difference spectrum of purified supercomplex after size exclusion chromatography (see panel A) in 100 mM Tris-HCl pH 7.5, 100 mM NaCl, 2 mM $MgSO_4$, 0.05% DDM. Peaks were fitted using Origin Pro 2016 (OriginLab corporation) showing an estimated ratio of approx. 1:1:1 of hemes *a*:*b*:*c* (1) (**C**) BN-PAGE of the purified supercomplex (SC peak in panel A) showing the presence of $III_2IV_2$ supercomplexes, confirmed using mass spectrometry. (**D**) SDS-PAGE of the purified supercomplex (SC peak in panel A). Bands were assigned according to size and (1).

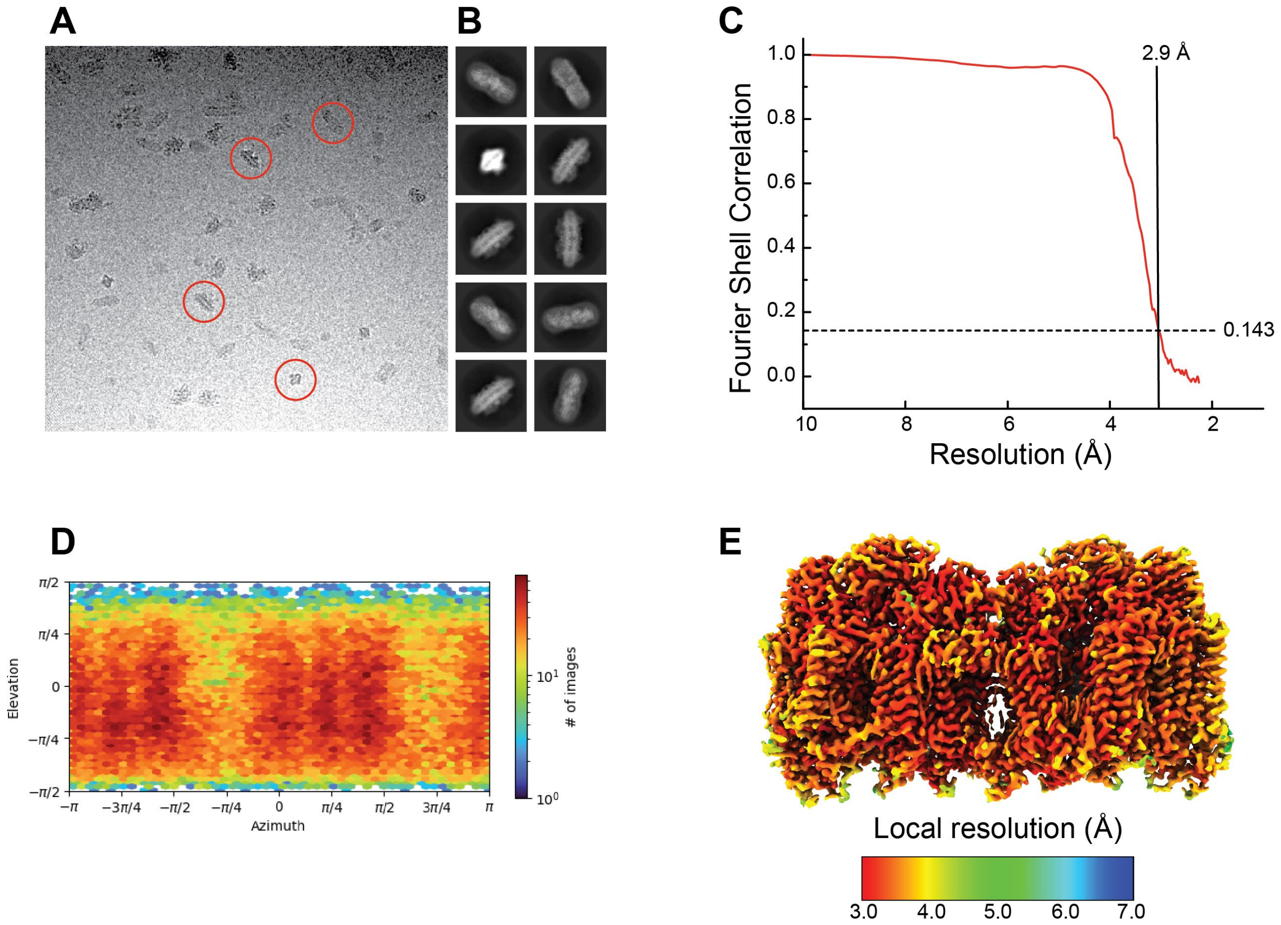

**Figure S2**. **Cryo-EM map validation.** (**A**) Example micrograph (**B**) 2D class averages (**C**) Fourier shell correlation (FSC) curve after refinement, corrected for the effects of masking (**D**) Viewing direction distribution for particle images (E) local resolution map.

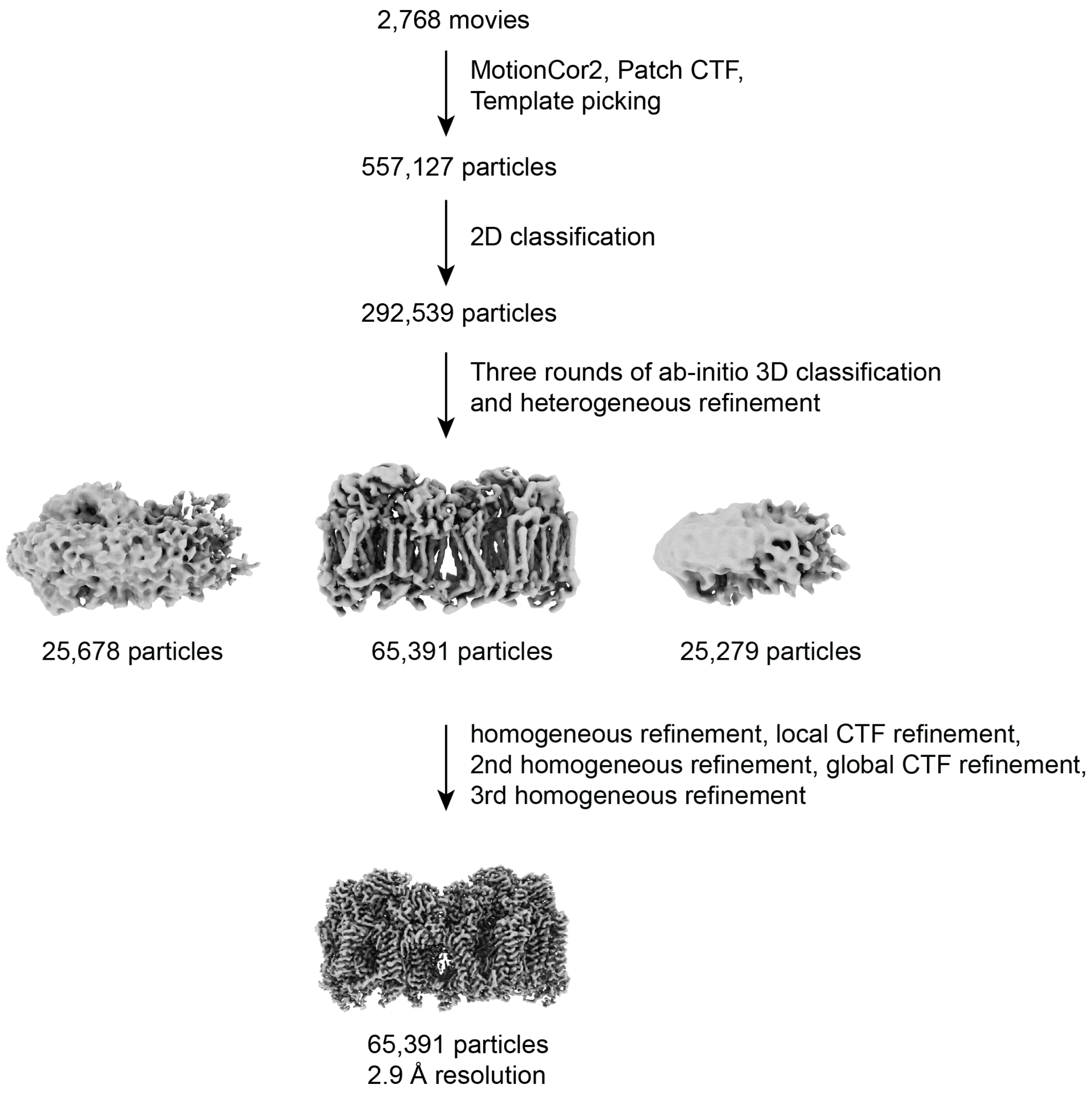

**Figure S3. Workflow for cryo-EM image analysis**

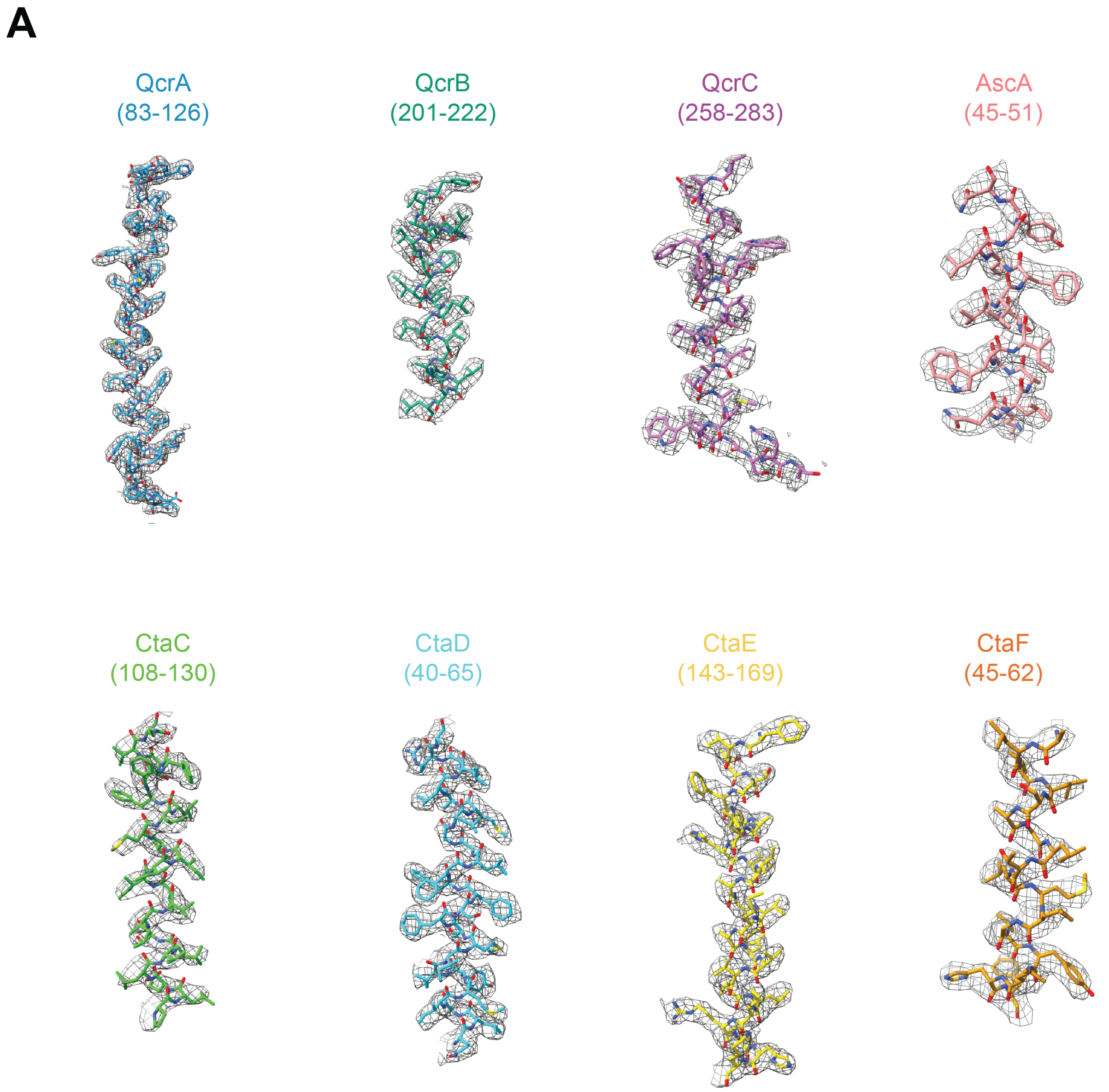

A
QcrA
(83-126)
QcrB
(201-222)
QcrC
(258-283)
AscA
(45-51)
CtaC
(108-130)
CtaD
(40-65)
CtaE
(143-169)
CtaF
(45-62)

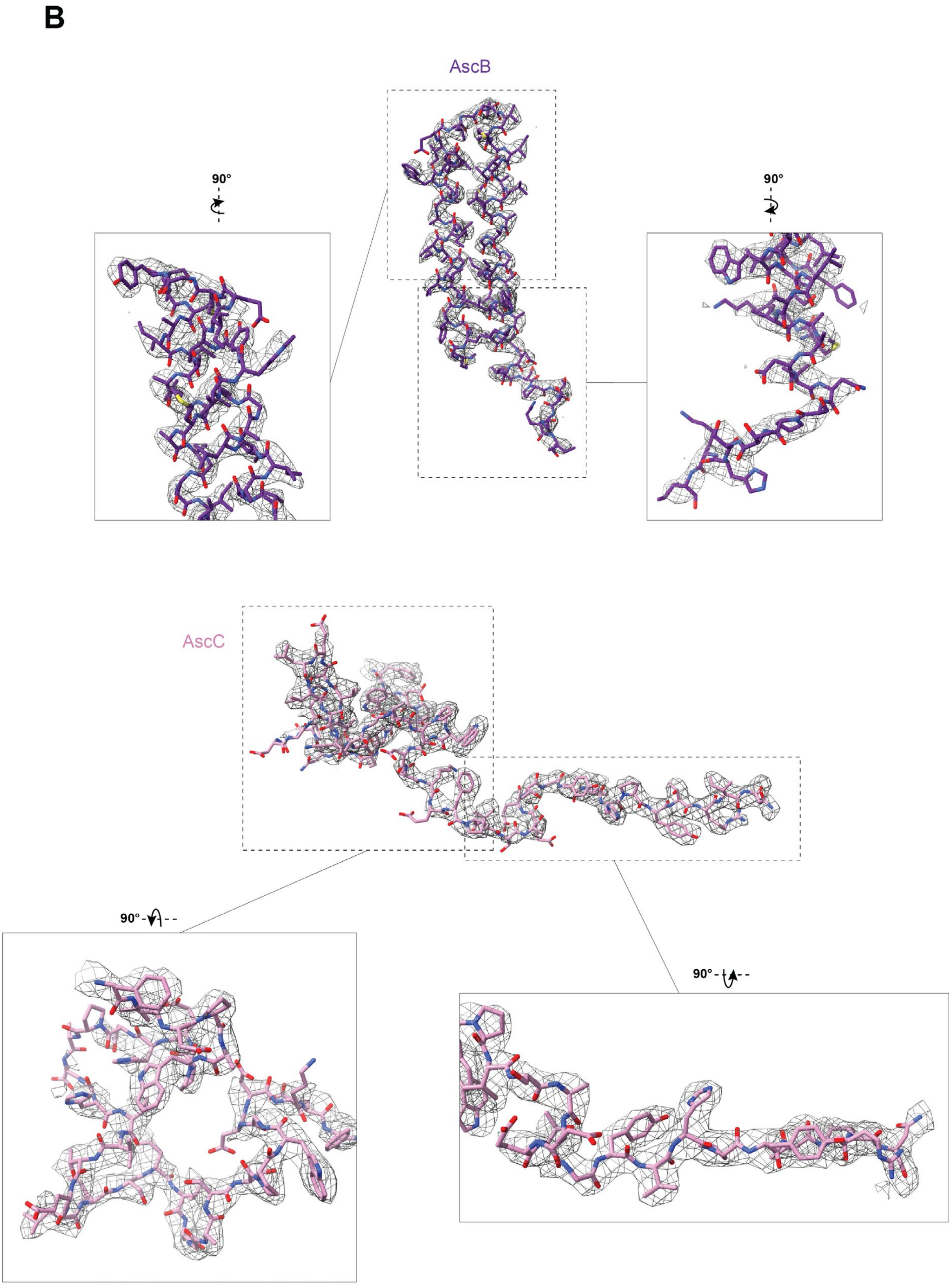

**Figure S4. Models and maps.** (**A**) Representative atomic model and experimental map for selected regions of the supercomplex. (**B**) Atomic model and experimental map of AscB and AscC.

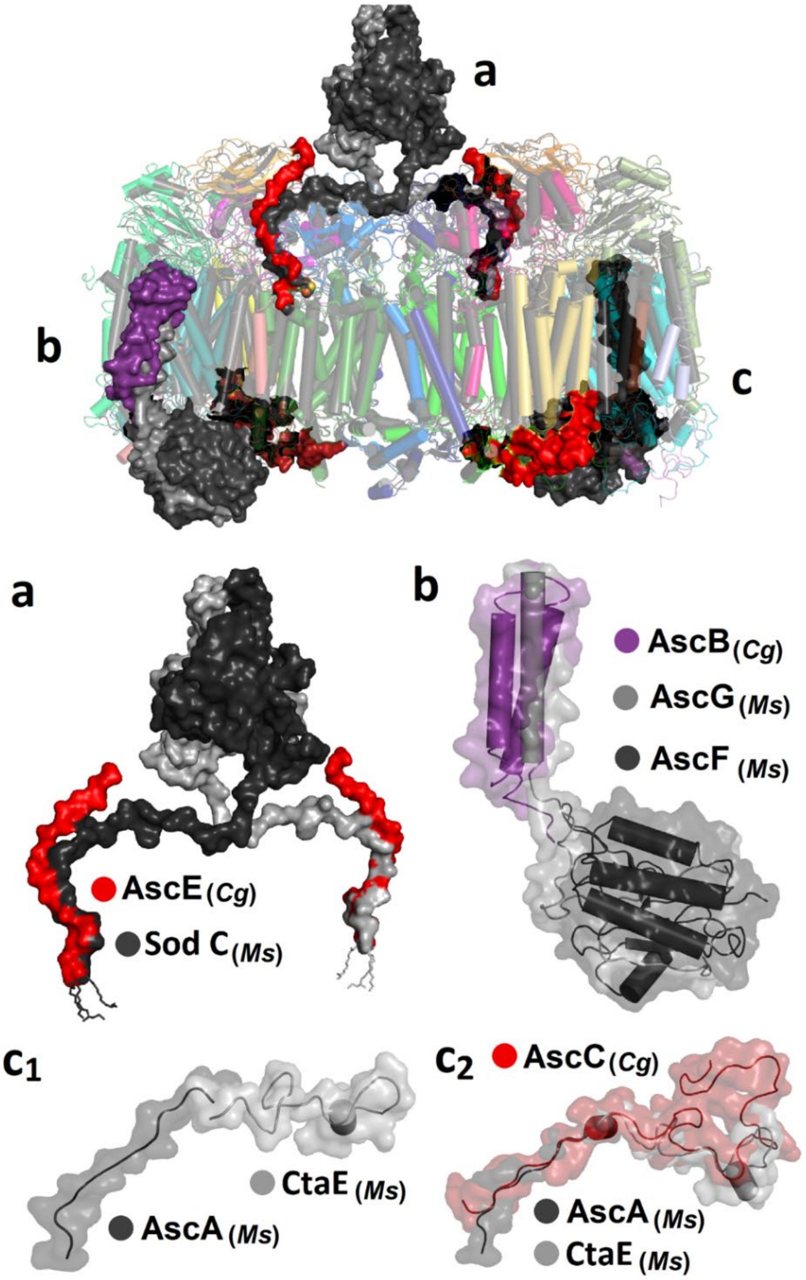

**Figure S5. Comparison of accessory subunits of the *C. glutamicum* and *M. smegmatis* supercomplexes.** The segments are indicated with letters a-c and shown in detail in the lower part of the figure. The transmembrane subunit AscB (*C. glutamicum*) overlaps in space with AscG (*M. smegmatis*, earlier MSMEG_4693). The *M. smegmatis* subunit AscF was earlier named MSMEG_4692.

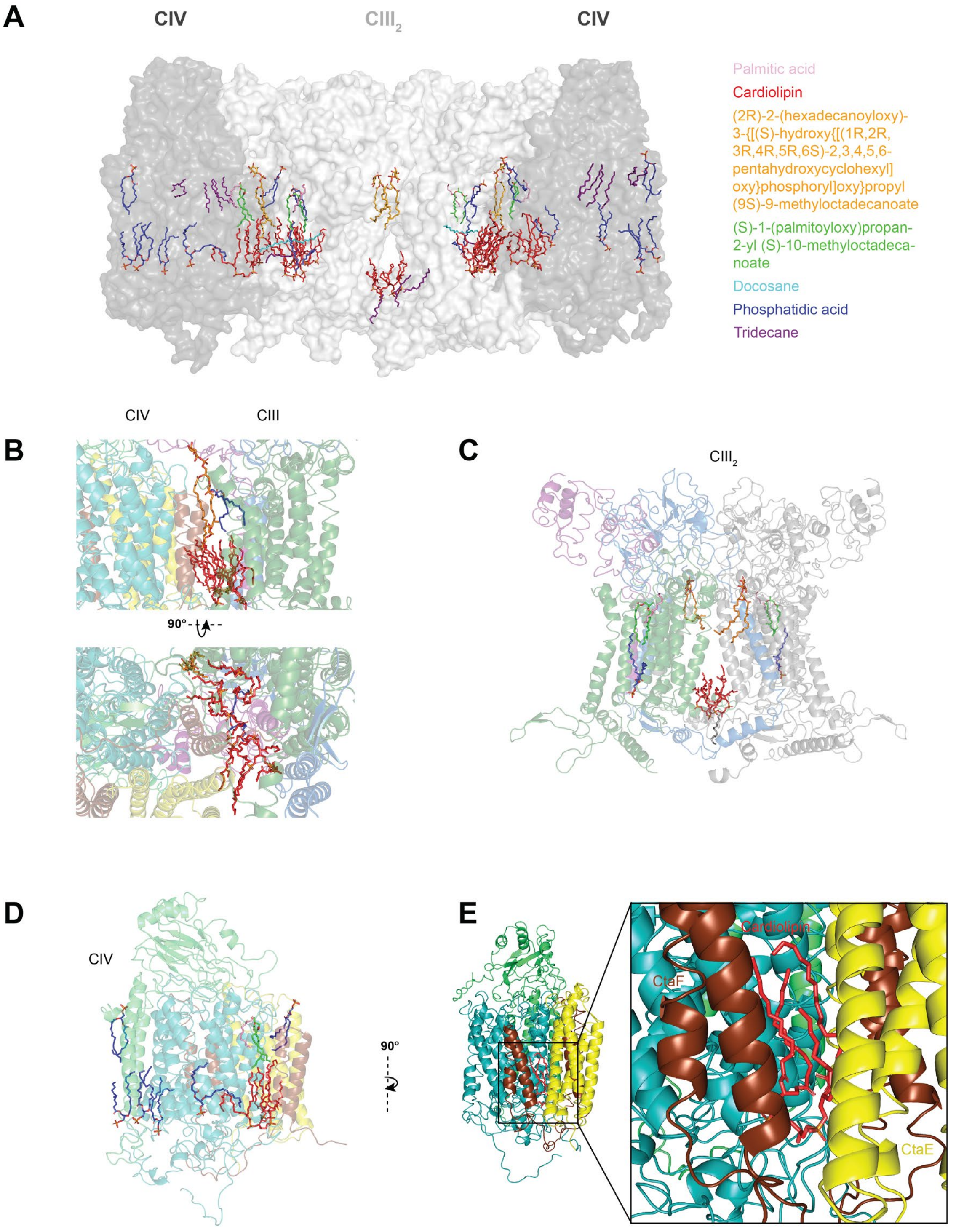

**Figure S6. Lipids in the $CIII_2CIV_2$ supercomplex**. (**A**) Density was attributed to 61 lipid molecules. Unidentifiable lipids were modeled as hydrocarbon chains. Lipids at the CIII-CIV interface (**B**), in $CIII_2$ (**C**) and in CIV (**D**). (**E**) Cardiolipin in a cavity defined by subunits CtaE and CtaF, suggested to be used for $O_2$ diffusion to the CIV catalytic site.

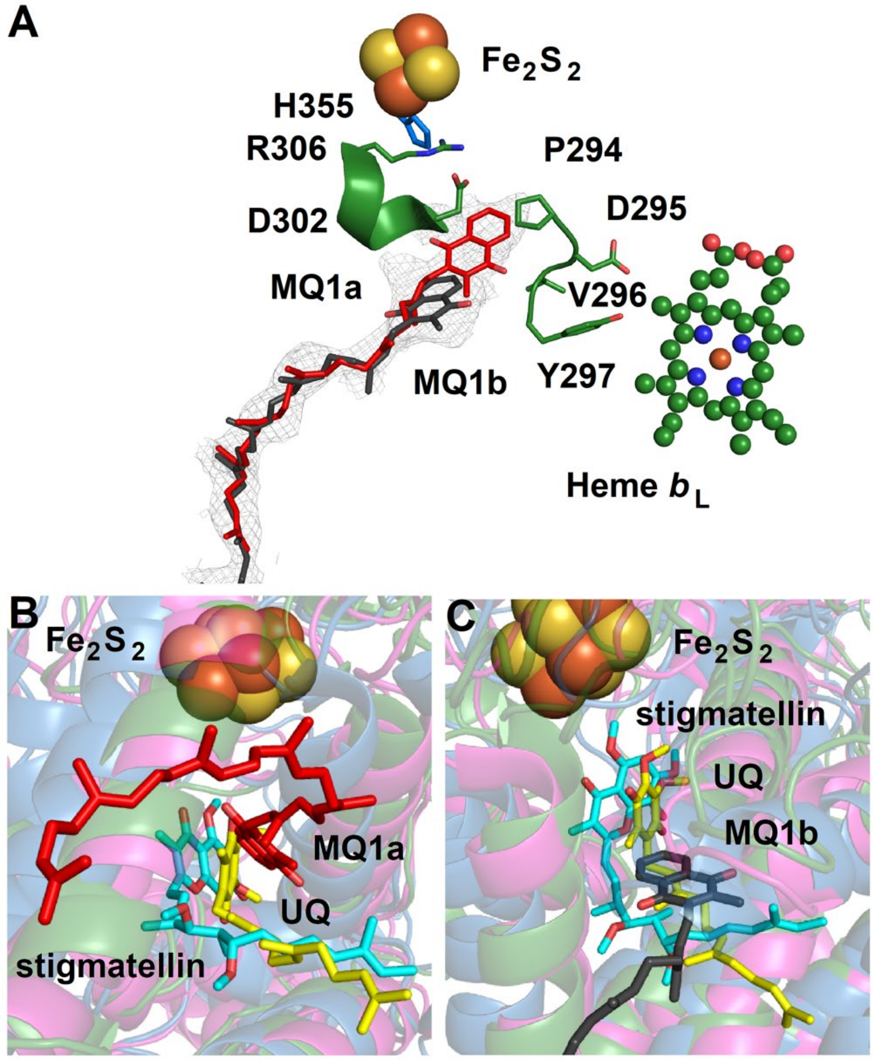

**Figure S7. The $Q_P$ site.** (**A**) The MQ density in *C. glutamicum* CIII overlaid with the MQ1a (colored in red) and MQ1b (colored in grey) sites in *C. glutamicum* and *M. smegmatis* CIII, respectively. (**B,C**) Superposition of *C. glutamicum* CIII (green helices) with MQ bound in sites MQ1a (**B**) or MQ1b (**C**) and UQ in *Ovis aries* (magenta helices, PDB: 6Q9E), and stigmatellin in *Rhodobacter capsulatus* (blue helices, PDB: 1ZRT).

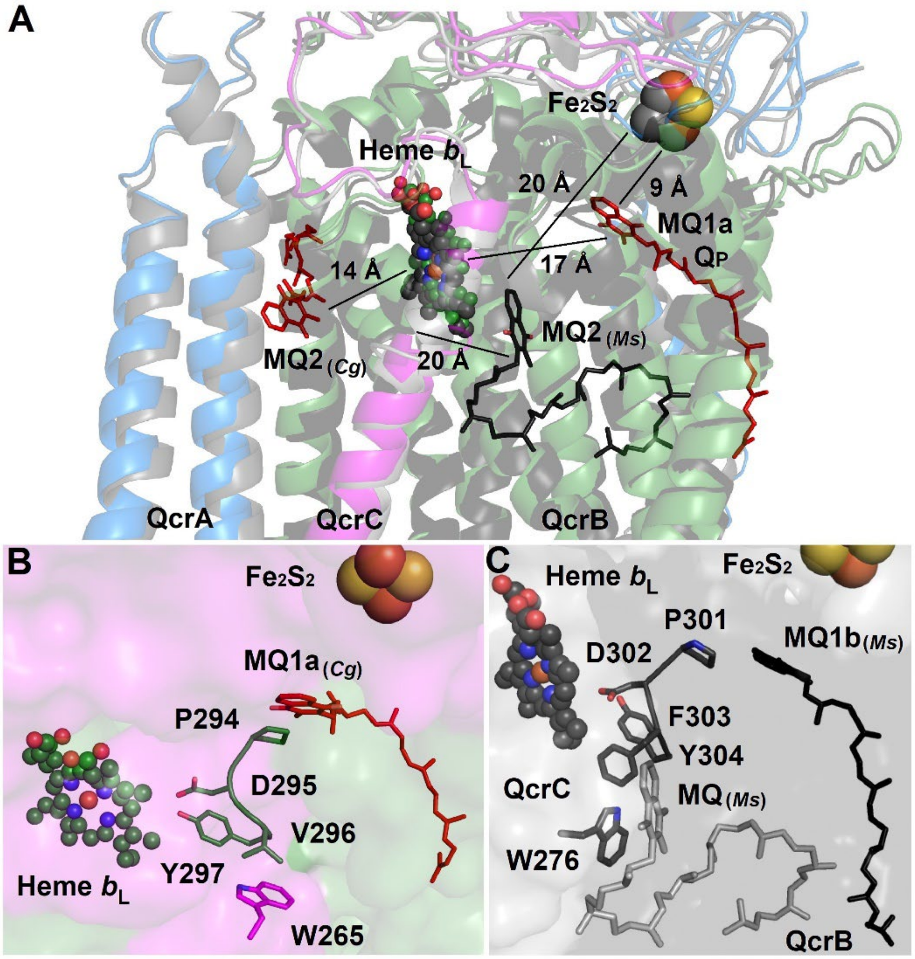

**Figure S8. MQ2 in CIII of *C. glutamicum* and *M. smegmatis*, respectively.** (**A**) The position of MQ2 in *M. smegmatis* CIII (*Ms* in black) is overlaid on the structure of *C. glutamicum* CIII. (**B**) Structure of a segment of *C. glutamicum* CIII where MQ2 is found in *M. smegmatis.* In *C. glutamicum*, binding of MQ in this location is obstructed by W265, which adopts a different position than in *M. smegmatis* due to the presence of a smaller V297 in the conserved Q-binding loop (PDVY). MQ1a is shown for reference. (**C**) The *M. smegmatis* MQ2 binding site ($MQ_{(Ms)}$) and the conserved Q-binding loop (PDFY). MQ1b is shown for reference.

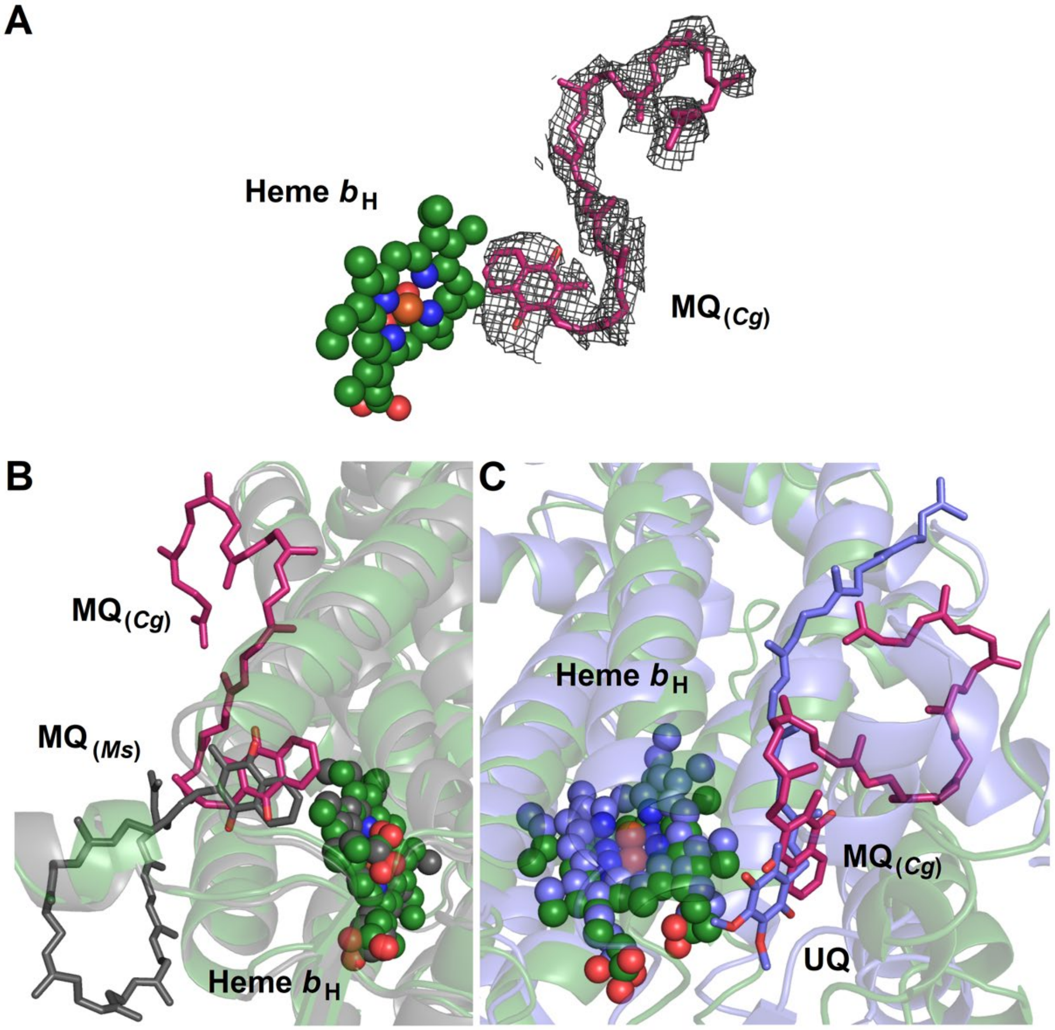

**Figure S9. The $Q_N$ site** (**A**) Density map (black) of MQ bound in $Q_N$ site of *C. glutamicum* CIII. (**B**) Superposition of the $Q_N$ sites of *C. glutamicum* (green helices) and *M. smegmatis* (grey helices; PDB: 6HWH) CIII with bound MQ (pink for *C. glutamicum* and grey for *M. smegmatis*). (**C**) Superposition of the $Q_N$ sites of *C. glutamicum* (green helices) and *S. cerevisiae* (purple helices, PDB: 4PD4) CIII with bound MQ (pink) or UQ (purple), respectively.

**Table S1. Identifiers for proteins of the *C. glutamicum* respiratory supercomplex.**

| Protein | Function | Cg annotation | NCgl annotation | Cgl annotation | Niebisch and Bott (2) |
|---|---|---|---|---|---|
| QcrA | Rieske FeS protein of complex III | Cg2404 | NCgl2110 | Cgl2190 | |
| QcrB | Cytochrome *b* of complex III | Cg2403 | NCgl2109 | Cgl2189 | |
| QcrC | Diheme cytochrome *cc* of complex III | Cg2405 | NCgl2111 | Cgl2191 | |
| CtaD | Subunit I of complex IV | Cg2780 | NCgl2437 | Cgl2523 | |
| CtaC | Subunit II of complex IV | Cg2409 | NCgl2115 | Cgl2195 | |
| CtaE | Subunit III of complex IV | Cg2406 | NCgl2112 | Cgl2192 | |
| CtaF | Subunit IV of complex IV | Cg2408 | NCgl2114 | Cgl2194 | |
| LpqE | Accessory protein | Cg2949 | NCgl2574 | Cgl2664 | P29 |
| AscA (PRSAF1) | Accessory protein | Cg2211 | NCgl1941 | Cg2211 | P20 |
| AscB | Accessory protein | Cg0775 | NCgl0643 | Cgl0673 | not detected |
| AscC | Accessory protein | Cg0935 | NCgl0784 | Cgl0818 | not detected |
| AscD | Accessory protein | unknown | unknown | unknown | not detected |
| AscE | Accessory protein | unknown | unknown | unknown | not detected |

**Table S2. Cryo-EM data collection, refinement and validation statistics**

| | |
|---|---|
| **Data collection and processing** | |
| Magnification | 130,000 |
| Voltage (kV) | 300 |
| Electron exposure ($e^-/Å^2$) | 55 |
| Defocus range (μm) | -1.0 to -2.4 |
| Pixel size (Å) | 1.06 |
| Symmetry imposed | None (C1) |
| Initial particle images (number) | 557,127 |
| Final particle images (number) | 65,391 |
| Map resolution (Å) | 3.0 |
| FSC threshold | 0.143 |
| Map resolution range (Å) | 2.4-4.3 |
| **Refinement** | |
| CC (mask) | 0,87 |
| Resolution estimates (Å) | |
| d 99 | |
| Masked | 3.2 |
| Unmasked | 3.2 |
| d FSC model, 0/0.143/0.5 (Å) | |
| Masked | 2.4/2.9/3.1 |
| Unmasked | 2.4/2.9/3.1 |
| Model composition | |
| Nonhydrogen atoms | 47731 |
| Protein residues | 5813 |
| Ligands | 92 |
| B factors ($Å^2$) | |
| Protein residues | 53.9 |
| Ligand | 87.0 |
| **Validation** | |
| MolProbity score | 2.2 |
| Root mean square deviations | |
| Bond lengths (Å) | 0.006 |
| Bond angles (°) | 1.38 |
| Clashscore | 12.3 |
| Poor rotamers (%) | 1 |
| Ramachandran plot | |
| Favored (%) | 93.2 |
| Allowed (%) | 6.5 |
| Disallowed (%) | 0.3 |
